# The reliability and accuracy of recombination inferred by Shapeit2 duoHMM on whole genome sequence

**DOI:** 10.64898/2026.05.06.723015

**Authors:** Samir Oubninte, Ingo Ruczinski, Lisa R. Yanek, Rasika Mathias, Alexandre Bureau

## Abstract

Few studies assessed the performance of population-based phasing combined with parental genotypes to infer recombination on whole genome sequence (WGS) data. In this study, our objective was to evaluate whether Shapeit2 duoHMM, a Hidden Markov Model using parental genotypes, infers recombination events reliably on WGS and with narrower intervals than SNP arrays. We based our analysis on the overlap between recombination events inferred by Merlin on SNP genotypes and Shapeit2 on WGS and SNP genotypes. We used a sample of 61 extended families from the GeneSTAR study with TopMED freeze 8 WGS on 580 sequenced subjects (60% of sample). Shapeit2 was run with a window size of 500 kilobases and 200 states on WGS. To mimic a SNP array, we extracted genotypes of 355,112 autosomal markers on the Illumina OmniExpress array. The number of recombination events per meiosis inferred by Shapeit2 on the WGS data (36.8) was aligned with the expected numbers over autosomes (35.7), although Merlin overestimated this number (115.0). 73% of Shapeit2 recombination events on WGS were detected by Merlin, a proportion rising to 91% when restricting to events also inferred by Shapeit2 on OmniExpress genotypes. Furthermore, Shapeit2 recombination intervals were narrower on WGS than OmniExpress genotypes (median of 4,530 bp vs. 49,458 bp). This suggests that Shapeit2 on WGS is a reliable and accurate method for inferring recombination events.

## 1 Introduction

Recombination is a fundamental biological process, essential for genetic diversity, evolution, DNA repair, and the prevention of harmful mutations, and it is crucial for the survival of sexually reproducing organisms (Baudat et al. 2013). Its study spans evolutionary biology, disease genetics, and biotechnology, offering insights into genetic variation and disease mechanisms while enabling applications in agriculture and medicine (Baudat et al. 2013; Pratto et al. 2014). Genotype data provides a comprehensive map of genetic variation within populations, serving as a foundation for understanding recombination mechanisms and the distribution of genetic diversity; with high-throughput sequencing technologies, researchers can now obtain vast amounts of genetic information from individuals, shedding light on DNA sequence variations across populations. In light of this, rigorous models of fundamental biological processes are essential, and central to these models is the precise localization of the recombination site, which is vital for determining the segment of DNA transmitted to the subsequent generation and identifying genes of interest. In this study, one of the proposed objectives is to accurately infer recombination events in families.

The precision of gene maps is predominantly determined by the number of actual recombination events present in the data. Numerous studies have demonstrated that the use of inaccurate gene maps during analysis can reduce power and introduce bias in the results, leading to significantly divergent results (He et al. 2011a). Precise recombination localization will enhance the accuracy of disease gene mapping studies (He et al. 2011a). The inference of recombination events remains a complex issue, becoming even more challenging when older generations are not genotyped. Few studies have evaluated the efficacy of population-based phasing in conjunction with parental genotypes to infer recombination on whole genome sequence (WGS) data (Halldorsson et al. 2019; O’Connell et al. 2014). Halldorsson et al. (2019) utilized SNP-chip-genotyped and whole-genome-sequenced Icelandic samples to identify parent-child pairs from a comprehensive genealogical database, determined crossover locations by analyzing haplotype phase transitions and further refined these locations using sequence-level data, but did not evaluate recombination inference performance. A limited number of methods exist for efficiently and accurately estimating recombination events for a large number of closely related markers from general pedigrees, and their performance has not been extensively assessed. Beyond O’Connell’s 2014 paper (O’Connell et al. 2014), there has been a lack of publications that directly examine the inference of recombination events. Several computational methods, such as Beagle (Browning and Browning 2011), have been proposed that allow these events to be indirectly inferred from available genotype data. These methods primarily target pedigrees of small complexity, with the objective of determining the most likely haplotypes (O’Connell et al. 2014; Abecasis et al. 2002; Zhang and Zhao 2006; Delaneau et al. 2012; Browning and Browning 2007; Blackburn et al. 2020). The switch error rate (SER) and various measures of genotype imputation accuracy serve as the main parameters to compare these methods (Huang et al. 2015; Mitt et al. 2017; Choi et al. 2018).

In our pursuit of accurately inferring recombination events, we draw comparisons between two widely used methods: Shapeit2-duoHMM (O’Connell et al. 2014) and Merlin (Abecasis et al. 2002). Merlin was developed for pedigree-based linkage analysis and haplotyping using marker maps (e.g., SNP arrays); here, it serves as a baseline for recombination event inference from genotyping arrays. Shapeit2-duoHMM leverages a hidden Markov model with parental genotypes to refine phasing and infer recombination events (O’Connell et al. 2014). While Shapeit2 and Merlin have outperformed other methods in nearly all scenarios (O’Connell et al. 2014; Abecasis et al. 2002; Zhang and Zhao 2006), Pulsar (Blackburn et al. 2020) achieves SER comparable to Shapeit2 on simulated data from small families where most individuals have been sequenced. To our knowledge, no publication has been identified that examines the recombination events inferred on the whole Genome Sequence (WGS) by Shapeit2-duoHMM and directly assesses the reliability and accuracy of the resulting recombination intervals.

The aim of this paper is demonstrating the efficacy of Shapeit2-duoHMM (O’Connell et al. 2014) in inferring recombination events from WGS in comparison with the performance of Merlin (Abecasis et al. 2002), particularly focusing on the precision of the inferred intervals compared to those obtained from Single Nucleotide Polymorphism (SNP) arrays. The analysis is based on the overlap between recombination events inferred by Merlin from genotype arrays and those inferred by Shapeit2-duoHMM from both genotype arrays and WGS. This comparative study offers an evaluation of the efficacy of these methods in deducing recombination events. Furthermore, it will serve as a benchmark, enabling us to establish a gold standard reference for less informative setups.

## 2 Materials and Methods

This section presents the dataset used in our study, describes the methodological framework used to infer recombination events with Shapeit2-duoHMM and Merlin, and details the strategy for their comparative analysis. Prior to analysis, we implemented rigorous quality control measures to ensure the reliability and accuracy of our findings.

### 2.1 Study Data

The study utilizes a dataset from the GeneSTAR (Genetic Study of Atherosclerosis Risk) study, a longitudinal cohort of families identified from probands with early-onset cardiovascular disease (National Center for Biotechnology Information (NCBI)), incorporating the TopMED freeze 8 whole genome sequencing data. Multiple sequencing centers performed the WGS. Sequencing was conducted to a median depth of 30X using DNA extracted from blood, employing PCR-free library construction and Illumina HiSeq X technology. This dataset includes 952 individuals from 61 extended families, comprising 351 founders and 601 non-founders. Fifty-one of these families span three generations, while the remaining 10 families cover two generations (refer to Table 1 for more descriptive statistics). The dataset features a diverse representation, including European American (EA) and African American families (AA) (Table 2), each with at least 8 sequenced members. In the GeneSTAR study, participant ancestry is typically determined through a combination of self-reporting and genetic analysis (Qayyum et al. 2015). Written informed consent was obtained from all participants, and the study was approved by the Johns Hopkins Institutional Review Board (IRB00143190).

**Table 1:** Descriptive Statistics for Family Composition.

|  | N | Minimum | Q1 | Median | Mean | Q3 | Maximum |
| --- | --- | --- | --- | --- | --- | --- | --- |
| Number of Individuals | 952 | 5 | 11 | 15 | 15.61 | 20 | 34 |
| Number of Females | 482 | 2 | 5 | 8 | 7.90 | 10 | 17 |
| Number of Males | 470 | 1 | 5 | 7 | 7.70 | 11 | 18 |

**Table 2:** Distribution of families and sequenced subjects across ethnic groups. The table summarizes the number of families and sequenced subjects for self-declared European-American and African-American groups, along with their respective percentages of sequenced subjects and the number of subjects in each group.The data shows that 61% of European-Americans and 59% of African-Americans were sequenced, resulting 60% of the total sample being sequenced.

|  | Number<br>families | Number of<br>of sequenced subjects | % of sequenced<br>within group | Total number<br>of subjects |
| --- | --- | --- | --- | --- |
| European-American | 38 | 338 | 61% | 558 |
| African-American | 23 | 231 | 59% | 394 |
| **Total** | **61** | **569** | **60%** | **952** |

In this study, we focused on retaining single nucleotide variants. Given the inconsistencies in the literature on WGS data quality control (Huang et al. 2015; Mitt et al. 2017; Choi et al. 2018), we defined a specific procedure for our data. Initial steps involved removing monomorphic sites and utilizing Plink to scrutinize Hardy–Weinberg equilibrium, detect Mendelian inconsistencies, and blanking implicated genotypes. Variants were excluded from the WGS data if they lacked a PASS status, exhibited a missingness rate greater than 5%, had a Hardy–Weinberg equilibrium *P* -value below 1 × 10^*−*6^, or were flagged as potential genotyping errors with a probability exceeding 0.95 (see sub-section 2.2). This approach aligns with methodologies discussed by Huang et al. (2015); Mitt et al. (2017). At the end of this process, 45,859,708 variants were remaining.

Given that our comparative study involved examining recombination events inferred from SNP arrays, we extracted genotypes from autosomal markers present on the Illumina OmniExpress array within the WGS dataset to mimic genotypes from an SNP array. Markers were extracted from bim files for the Illumina OmniExpress array, with physical positions lifted over from hg19 to hg38 using the liftOver function of the R package rtracklayer with hg19ToHg38.over.chain. Variants in the WGS were extracted if they matched these positions and either alleles in the original OmniExpress bim file or alleles obtained after flipping strands to conform to the Haplotype Reference Consortium strands using the HRC-1000G-check-bim.pl script from www.well.ox.ac.uk/∼wrayner/tools.

This procedure matched 355,112 autosomal markers.

Merlin encounters limitations when managing large families. In our case, 30 out of 61 families with three generations were too large for Merlin’s analysis (refer to Supplementary Table S1A for detailed information on these families). Consequently, we decided to split each family into two subfamilies, ensuring an even distribution of full siblings. Custom code developed for this purpose automated the process, except for eight large families (e.g., families with inbreeding) necessitating manual splitting. This method guaranteed that each subfamily comprised roughly half of the children from the original nuclear family. For a visual representation, refer to the pedigree in Supplementary Figure S1, which illustrates the family structure before and after the division. Supplementary Table S1 delineates the composition of subfamilies. These updated (split) families were used for all methods applied in this study.

### 2.2 Inferring recombination by Shapeit2-duoHmm

To infer recombination events, we adhered to the steps recommended by Shapeit2 duoHMM authors. The phasing process for each autosome was conducted in two stages. Initially, we filtered out variants flagged for potential genotyping errors by the duoHMM algorithm, which identifies incongruent genotypes in duo (parent-child) and trio (parent-parent-child) configurations (see Supplementary Methods). The cleaned data then underwent a second phasing run, yielding SHAPEIT2’s diploid graph representations of haplotype structure for downstream analyses. Phasing was the most time-consuming step, requiring between 382.5 hours for chromosome 21 and 562.8 hours for chromosome 1 using eight parallel threads. To integrate over phase uncertainty in crossover detection, we sampled haplotypes from Shapeit2’s diploid graph using ten distinct random seeds. For each sampled haplotype set, recombination probabilities were calculated using duoHMM, and the resulting maps were averaged. The final output detailed the parent and child involved in each meiosis, the regions where crossovers may have occurred (flanked by heterozygote sites on the parents), and the probability of a crossover event occurring. To accurately phase the whole-genome sequencing data, we employed Shapeit2 duoHMM with a window size of 500 kilobases and increased the state space from 100 to 200, as a larger state space improves accuracy, albeit with a linear increase in computing time. Despite implementing all recommended preprocessing steps to prevent bugs, Shapeit2 failed to converge during the initial run on chromosome 9 and terminated without generating an explicit error message. We addressed this issue by splitting chromosome 9 into its p and q arms which allowed the phasing and subsequent recombination inference to be completed successfully on each arm.

### 2.3 Inferring recombination From Merlin outputs

Merlin infers recombination by using sparse binary trees to represent gene flow patterns in pedigrees. These trees allow Merlin to efficiently calculate likelihoods and probabilities of different inheritance patterns, taking into account the recombination events that have occurred between markers. By analyzing the sequence of binary digits in the inheritance vectors, Merlin can identify the most likely paths of gene flow through the pedigree, considering the recombination events that separate markers. This approach enables Merlin to infer recombination events and estimate haplotypes, lod scores, and pairwise IBD probabilities (Abecasis et al. 2002). Unlike Shapeit2-duoHMM, Merlin’s standard output for gene flow (the flow file) provides the single most likely inheritance pattern rather than a posterior probability for each identified recombination event.

The flow file generated by Merlin provides a detailed record of genetic inheritance within a pedigree and tracks the descent of haplotypes across the pedigree, highlighting recombination events and inheritance patterns. In this file, each row corresponds to a single haplotype; therefore, two consecutive rows represent one individual (one maternal and one paternal haplotype). Using a unique identifier assigned to each individual—repeated on both haplotype rows—the file facilitates tracing the descent of founder alleles through the pedigree and identifying IBD relationships between individuals. Within each haplotype row, alleles are arranged sequentially according to their chromosomal positions, and non-founders display the alleles inherited from their parents (see Supplementary Figure S4). This structure enables clear lineage tracing and the detection of recombination events.

Inferring recombination events involves identifying the parent-offspring pairs between which the recombination occurs, as well as determining the boundaries of the recombination interval. We detect transitions in the offspring’s haplotype relative to the parent, which indicate the occurrence of a recombination event. The interval is then defined by the two flanking markers where the relevant parent is heterozygous, similar to the approach described by O’Connell et al. (2014).

### 2.4 Analysis of recombination events

In this study, we compared the number of recombination events and the number of events per meiosis to the expected autosomal counts. We also contrasted Shapeit2-duoHMM recombination intervals in WGS data with those inferred from OmniExpress genotypes using both Shapeit2-duoHMM and Merlin (see flow in Figure 1). The expected counts were derived from the number of parent–child pairs and genetic distances reported for the European population in Table 3 of He et al. (2011b). We conducted a concordance analysis by counting overlaps between intervals produced at loci where recombination events were likely to have occurred. Confidence intervals were calculated using the Wald method for proportions, leveraging the inter-family distribution, as each family represents an independent entity. Finally, we performed an ethnic diversity analysis involving European American and African American families from the GeneSTAR sample.

**Figure 1:**
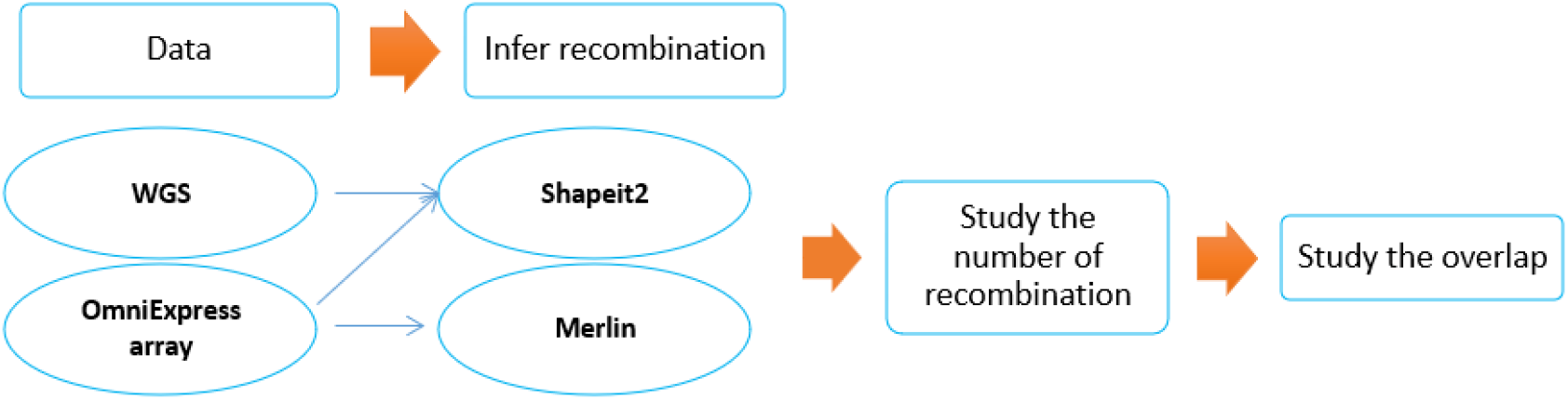
Recombination events analysis workflow: This diagram delineates the process involved in our genetic recombination analysis: starting with data processing, followed by the inference of recombination events and ending by examining their number and overlap.

## 3 Results

The analysis of recombination events inferred by Shapeit2 duoHMM on WGS data revealed the following results:

### 3.1 Recombination Event Detection in Parent-Child Pairs

We focused on identifying recombination events using three distinct Method-dataset combinations: Shapeit2-duoHMM on whole genome sequences (Shapeit2_WGS), Shapeit2-duoHMM on the Illumina OmniExpress array (Shapeit2_OE), and Merlin on the Illumina OmniExpress array (Merlin_OE), among 295 parent-child pairs common to the three settings. Recombination events inferred by Shapeit2-duoHMM were filtered with a posterior probability threshold of 0.5. Merlin does not provide an event-level posterior probability or equivalent confidence score, no such filtering could be applied. Subsequently, we compared these inferred events with the expected number of recombination events for each chromosome, calculated based on genetic distance.

As shown in Figure 2 and Supplementary Table S2, the number of recombination events inferred by Shapeit2-duoHMM in both WGS and SNP array data roughly matched the expected numbers over autosomes. A slight overestimation of recombination events inferred from WGS was observed for chromosome 9, which is likely related to processing the p and q arms separately. However, this localized excess has a negligible impact on the overall genomic results. Shapeit2-duoHMM identified 36.8 events in WGS data and 35.8 events in SNP array data. Conversely, Merlin significantly overestimated the number of recombination events, detecting 115.0 events compared to the expected 35.7 (see Supplementary Table S2).

**Figure 2:**
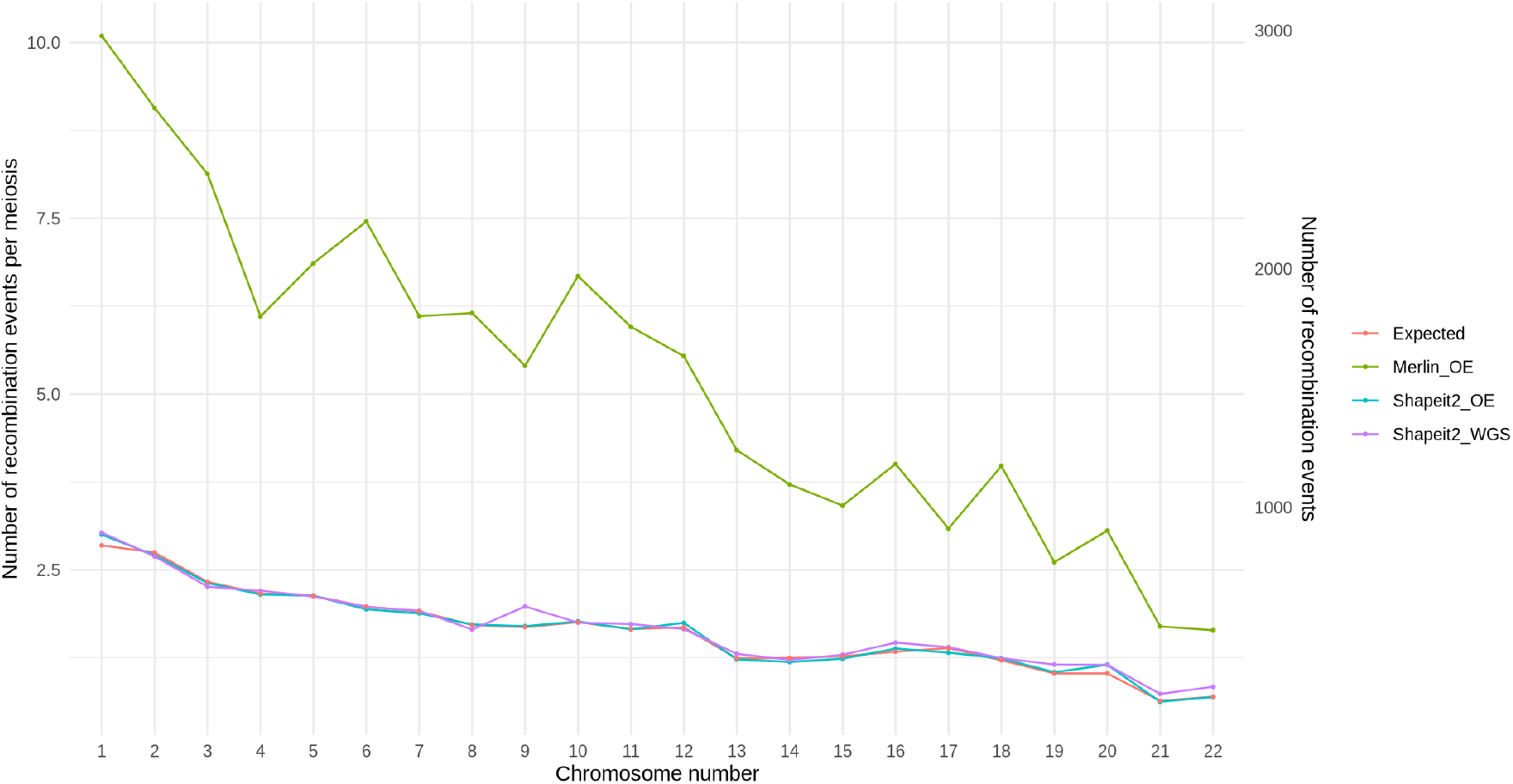
Distribution of Recombination Events Across Chromosomes. This figure illustrates the number of recombination events across chromosomes compared to the expected values. The ‘Merlin_OE’ represents OmniExpress genotypes data processed using the Merlin software package, while ‘Shapeit2_OE’ represents data processed with the Shapeit2 applied software package to OmniExpress data, and ‘Shapeit2_WGS’ represents data processed with Shapeit2 applied to complete sequence data.

### 3.2 Concordance of Recombination Events

Concordance for Shapeit2, as defined by O’Connell et al. (2014), was calculated as the proportion of Shapeit2 crossover events that overlap with a Merlin crossover event. Across all 61 families, 73.2% (95% CI: 72.4% to 74.0%) of Shapeit2 recombination events inferred from WGS data were also detected by Merlin. This proportion increased to 90.9% (95% CI: 90.2% to 91.5%) when restricting to events inferred by Shapeit2-duoHMM on both WGS and OmniExpress genotypes (Figure 3). Regarding the OmniExpress SNP chip data, 82.2% (95% CI: 81.4% to 82.9%) of the recombination events detected by Shapeit2-duoHMM overlapped with those identified by Merlin. These results are consistent with the rates of 80% to 89% previously reported by the Shapeit2-duoHMM authors (O’Connell et al. 2014). It is observed that the Shapeit2 intervals inferred from SNP chip markers are larger compared to those from WGS data (see Figure 4 and Supplementary Figure S2), which increases the likelihood of overlapping with Merlin-inferred recombination events. Overall, the findings highlight a moderate concordance between the Merlin and Shapeit2 methodologies across chromosomes.

**Figure 3:**
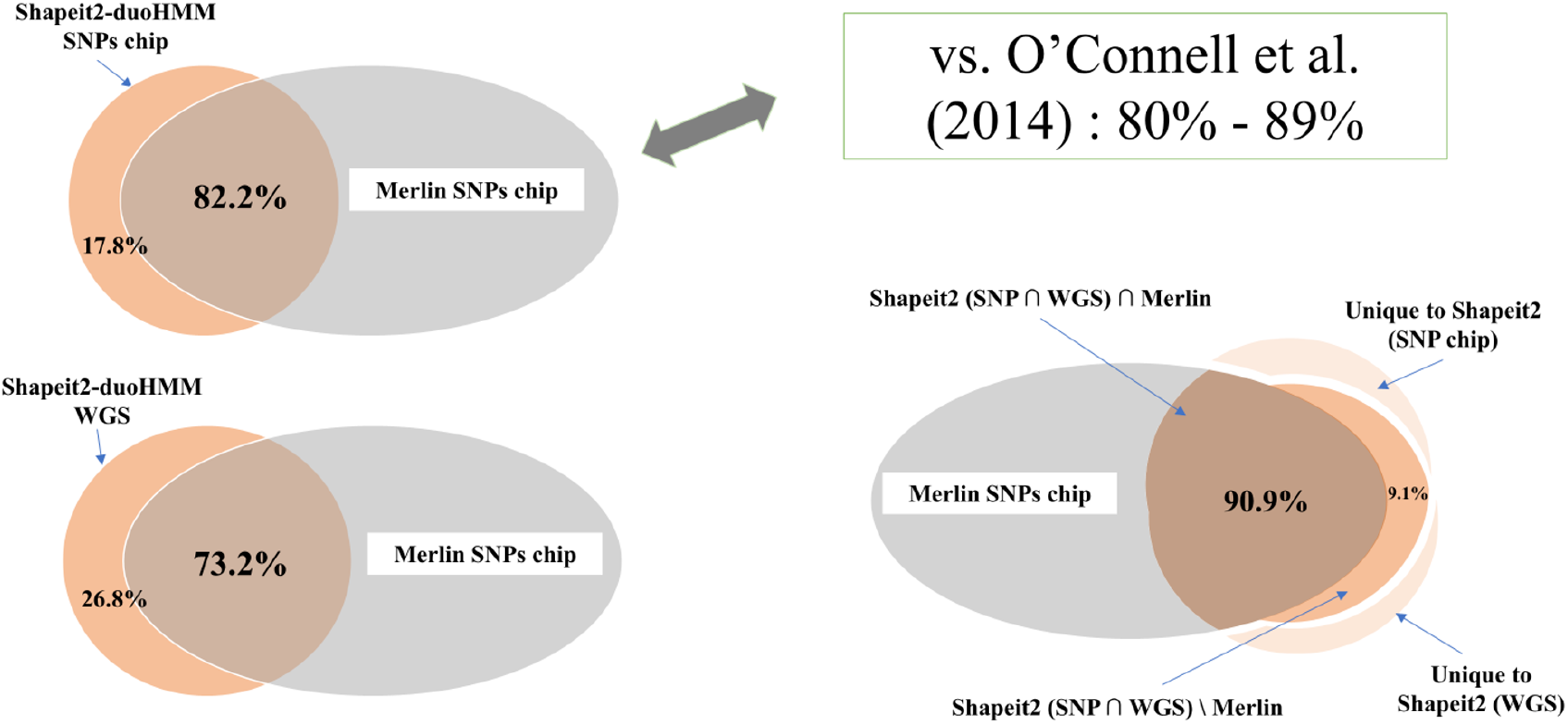
Detection by Merlin of Shapeit2 recombination events inferred on different genotype densities. top-left) Proportion of Shapeit2 recombination on SNPs chip detected by Merlin, bottom-left) Proportion of Shapeit2 recombination on the WGS detected by Merlin, bottom-right) Proportion of Shapeit2 recombination (in both SNP chip & WGS) detected by Merlin

**Figure 4:**
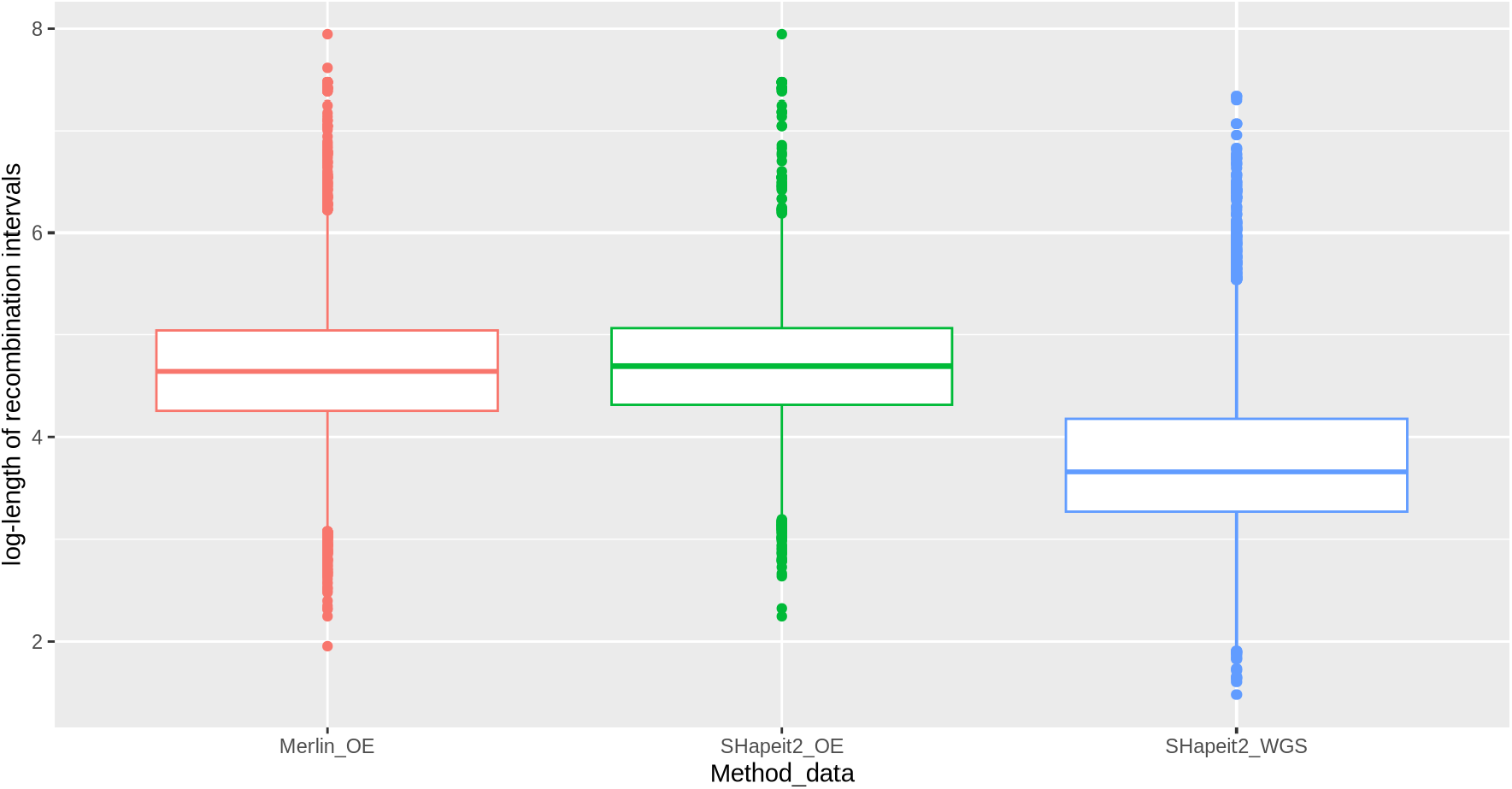
This figure illustrates the distribution of recombination event interval lengths on a log scale across chromosomes. The ‘Merlin_OE’ dataset represents OmniExpress genotype data processed using the Merlin software package, while ‘Shapeit2_OE’ represents data processed with the Shapeit2 software package applied to OmniExpress data, and ‘Shapeit2_WGS’ represents data processed with Shapeit2 applied to WGS data.

### 3.3 Precision of inferred recombination events

Figure 4 illustrates the distribution of interval lengths for the identified recombination events across autosomes. The precision of these intervals is primarily constrained by the density of informative (heterozygous) markers flanking each crossover. As expected, Shapeit2-duoHMM recombination events were localized within significantly narrower intervals on WGS data compared to OmniExpress genotypes, with median lengths of 5,530 bp and 49,458 bp, respectively. This marked difference reflects the vastly higher marker density of WGS, which allows for a more refined localization of the recombination event.

Furthermore, the intervals inferred by Shapeit2-duoHMM on SNP array data were comparable to those determined by Merlin, with median lengths of 49,458 bp and 44,310 bp, respectively (see Supplementary Table S3 and Supplementary Figure S2 for further details). Despite the substantially larger number of recombination events inferred by Merlin, the corresponding interval length distribution remains similar to that obtained with Shapeit2-duoHMM on the same SNP array data. This confirms that interval length is a function of the underlying marker density and parental heterozygosity—representing mapping resolution—rather than the specific inference algorithm employed.

### 3.4 Genetic Diversity and Recombination Events

The samples analyzed by O’Connell et al. (2014) were either Europeans or Africans, whereas the GeneSTAR cohort includes individuals who self-identify as European American (EA) and African American (AA), as shown in Table 2. To assess the impact of this diversity, we conducted separate analyses for these groups using Shapeit2-duoHMM recombination inference on OmniExpress SNP array. The number of recombination events per meiosis inferred for the EA group (36.1) was close to the expected value (35.7) and to the estimate for the entire sample (35.8) (Supplementary Figure S3, top). In contrast, fewer recombination events per meiosis were inferred in the AA group (32.0) (Mann–Whitney U test, *W* = 60, *p* = 0.03637). This discrepancy reflects differences in the inferred data and may be influenced by using European genetic maps to infer recombination events. Consequently, employing African-specific genetic maps may yield more accurate expected values for the AA group. Supplementary Figure S3 (bottom) shows recombination event interval distributions on a log scale, indicating visual consistency across methods. The geometric mean of the intervals was 16% shorter for the AA group than for the EA group (Mann–Whitney U test, *W* = 50, *p* = 0.01045), indicating that crossovers in the AA group tend to occur closer together, potentially reflecting population-specific differences in recombination patterns.

## 4 Discussion

Shapeit2 and Merlin estimate the patterns of inheritance and recombination in genotype data using distinct approaches. Choi et al. (2018) demonstrated that integrating parental data into Shapeit2 significantly improved haplotype accuracy and reduced switch error rates, outperforming all other tested methods. Shapeit2 utilizes a Hidden Markov Model (HMM) framework, considering population genetic principles and linkage disequilibrium patterns combined with constraints from genotypes of parent-child duos to identify allele inheritance and likely recombination regions. In contrast, Merlin is a pedigree-based tool that relies on known relationships and Mendelian inheritance laws to infer expected genotypes and recombination events within pedigrees by examining allele transmission across generations.

Inferring reliable and accurate recombination events remains a topic of significant interest with great potential for genomic data analysis. To investigate this, we utilized the GeneSTAR cohort, which consists of related individuals who have been genotyped and sequenced, to evaluate the Shapeit2-duoHMM and Merlin approaches for inferring recombination events. Shapeit2-duoHMM proved to be both reliable and accurate in inferring recombination events from whole-genome sequence data. The number of inferred events aligned closely with chromosomal expectations. A substantial proportion of Shapeit2-duoHMM-inferred events were detected by Merlin, with this proportion improving when focusing on events inferred from both genotype and sequence data. Furthermore, recombination events identified uniquely by Shapeit2-duoHMM in WGS data may represent genuine crossovers missed by SNP arrays, highlighting the superior resolution of sequencing data. We found no evidence of systematic under- or overestimation in the Shapeit2-duoHMM approach. In contrast, Merlin tended to overestimate the number of events; a significant proportion of these excess events likely represent false positives (no threshold-based filtering could be applied to calibrate the number of inferred events on expectations and thus reduce false positives). Finally, the larger median size of the recombination intervals observed with OmniExpress genotypes suggests a broader range of uncertainty.

Separate analyses of European American and African American families revealed a 16% difference in the mean, which is sufficiently small to suggest that these methods are robust across diverse genetic backgrounds. While sex-specific differences in recombination rates are well-recognized and extensively documented (O’Connell et al. 2014; Kong et al. 2010; Sardell and Kirkpatrick 2020; Gruhn et al. 2013), our primary objective was to evaluate the performance of inference methods rather than to characterize sex-specific patterns. Implementing sex-stratified analyses would require substantially larger sample sizes to preserve statistical power and robustness. We therefore adopted a sex-pooled approach to maintain the integrity of comparisons and align with the broader scope of this study. This design supports general applicability across the dataset.

While Shapeit2-duoHMM and Merlin are widely used tools for genetic analysis, they come with certain limitations and challenges. Both tools can be computationally intensive, particularly when dealing with large datasets or complex pedigrees, necessitating substantial computational resources such as memory and processing power. They require specific input and output formats, making data preprocessing, post-processing and conversion necessary. This process can be time-consuming and may introduce errors. Our findings confirm the limitations of SNP arrays in accurately capturing recombination events and reinforce the superior value of WGS data in genetic research. This emphasizes the importance of using WGS data to achieve more accurate and detailed insights into genetic recombination.

O’Connell et al. (2014) demonstrated the effectiveness of Shapeit2-duoHMM in inferring accurate recombination events by comparing the number of recombination events. In this study, we build on O’Connell’s work by further validating the reliability and accuracy of the inferred recombination events through a comparison of their overlap with Merlin. Additionally, we examined the behavior of recombination event intervals and assessed the performance of Shapeit2-duoHMM on whole-genome sequencing data. Our findings show that recombination events are precisely localized by the narrower intervals obtained with Shapeit2-duoHMM on WGS data, enhancing our understanding of recombination dynamics and advancing population genetics research. Based on the results of our study, we recommend for WGS the strategy of O’Connell et al. (2014): estimating haplotypes with Shapeit2 to phase genotype data, followed by the application of duoHMM, to infer accurate recombination events.

## 5 Code and Data availabilities

- The custom code developed for this research is central to our conclusions and is made available to ensure transparency and reproducibility of our results. The code-base, including all relevant custom scripts, has been open-sourced under the MIT license and is accessible via our GitHub repository at [https://github.com/oubninte/Shapeit2-duoHMM_on_WGS]. We encourage users to review the license for any usage restrictions that may apply.
- GeneSTAR data are available through the database of Genotypes and Phenotypes (dbGaP), study accession phs001218.

## 6 Funding and Acknowledgements

- This work was made possible by funding from the Natural Sciences and Engineering Research Council of Canada (NSERC).
- We would like to thank all GeneSTAR families for participating in the study.
- The GeneSTAR study was supported in part by grants from the National Heart, Lung, and Blood Institute (HL112064, HL72518, and HL087698), the National Center for Research Resources (M01-RR000052), and by the Intramural Research Program of the National Human Genome Research Institute, National Institutes of Health.
- Molecular data for the Trans-Omics in Precision Medicine (TOPMed) program was supported by the National Heart, Lung and Blood Institute (NHLBI). Genome Sequencing for “NHLBI TOPMed: Genetic Studies of Atherosclerosis Risk” (phs001218) was performed at Psomagen (3R01HL112064-04S1), Illumina (R01HL112064), and Broad Genomics (HHSN268201500014C). Core support including centralized genomic read mapping and genotype calling, along with variant quality metrics and filtering were provided by the TOPMed Informatics Research Center (3R01HL-117626-02S1; contract HHSN268201800002I). Core support including phenotype harmonization, data management, sample-identity QC, and general program coordination were provided by the TOPMed Data Coordinating Center (R01HL-120393; U01HL-120393; contract HHSN268201800001I). The authors would like to acknowledge contributions from the in-vestigators of the NHLBI TOPMed Consortium (https://www.nhlbiwgs.org/topmed-banner-authorship).
- I. Ruczinski was supported by NIH (grants R01 GM083084 and R01 DE031855).
- R.A. Mathias was supported in part by the MOSAIC award from The Johns Hopkins University Provost’s Office.
- This research was supported in part by the Intramural Research Program of the National Institutes of Health (NIH). The contributions of the NIH author were made as part of their official duties as NIH federal employees, are in compliance with agency policy requirements, and are considered Works of the United States Government. However, the findings and conclusions presented in this paper are those of the author and do not necessarily reflect the views of the NIH or the U.S. Department of Health and Human Services.

