## Supplementary information for "The reliability and accuracy of recombination inferred by Shapeit2 duoHMM on whole genome sequence"

### Supplementary Methods:

**Genotype Error Detection:** By comparing observed genotypes with expected patterns based on population genetics principles and pedigree information. these algorithms can identify likely errors in the genotype data. To detect unlikely genotypes. scientists compare the likelihood of observed genotypes assuming two scenarios: one where markers are linked (close together) and another where they are unlinked (far apart) [1]. They calculate a ratio of these likelihoods. If a genotype fits well with its neighboring markers. the ratio should be small. However. if a genotype does not fit well. the ratio becomes large. indicating a potential error. In a typical situation without errors. the chance of seeing a large ratio is very low. To flag problematic genotypes. they set a high threshold for this ratio [1]. The error detection algorithm in Shapeit2 uses a Hidden Markov Model (HMM) framework to model expected inheritance and recombination patterns in a population. comparing them to observed genotype data across individuals and markers. Integrated as a post-processing tool. DuoHMM refines haplotypes by incorporating pedigree information. detecting recombination events. correcting phasing errors to align with pedigree structures. and identifying subtle inconsistencies that may indicate genotyping errors [2]. Similarly. Merlin utilizes an error detection algorithm within its pedigree-based analysis framework. examining genotype patterns within pedigrees and comparing them to expected inheritance patterns. Both Shapeit2 and Merlin consider various factors when detecting genotyping errors. including Mendelian inheritance laws. relationships between individuals in a pedigree. and genotype consistency across related individuals [1.2]

| A) BEFORE (30 families) | N | min | Q1 | median | mean | Q3 | max |
|---|---|---|---|---|---|---|---|
| **Number of Individuals** | 630 | 13 | 18 | 20 | 21.00 | 25 | 34 |
| **Number of Females** | 315 | 3 | 9 | 10 | 10.5 | 12 | 17 |
| **Number of Males** | 315 | 5 | 8 | 11 | 10.5 | 12 | 18 |

| B) AFTER (60 subfamilies) | N | min | Q1 | median | mean | Q3 | max |
|---|---|---|---|---|---|---|---|
| **Number of Observations** | 682 | 5 | 9 | 11 | 11.37 | 14 | 21 |
| **Number of Females** | 341 | 2 | 4 | 6 | 5.68 | 7 | 10 |
| **Number of Males** | 341 | 2 | 4 | 5 | 5.68 | 7 | 11 |

**Supplementary Table S1:** Pedigree statistics before (top) and after (bottom) splitting the families into subfamilies based on kinship information. Siblings (kinship = 0.250) within each family were separated into two groups to construct their genealogies. The observed differences in the number of individuals and observations between the two states are due to founders who are duplicated across some of the subfamilies. Two out of the 60 subfamilies spanning only two generations instead of three.

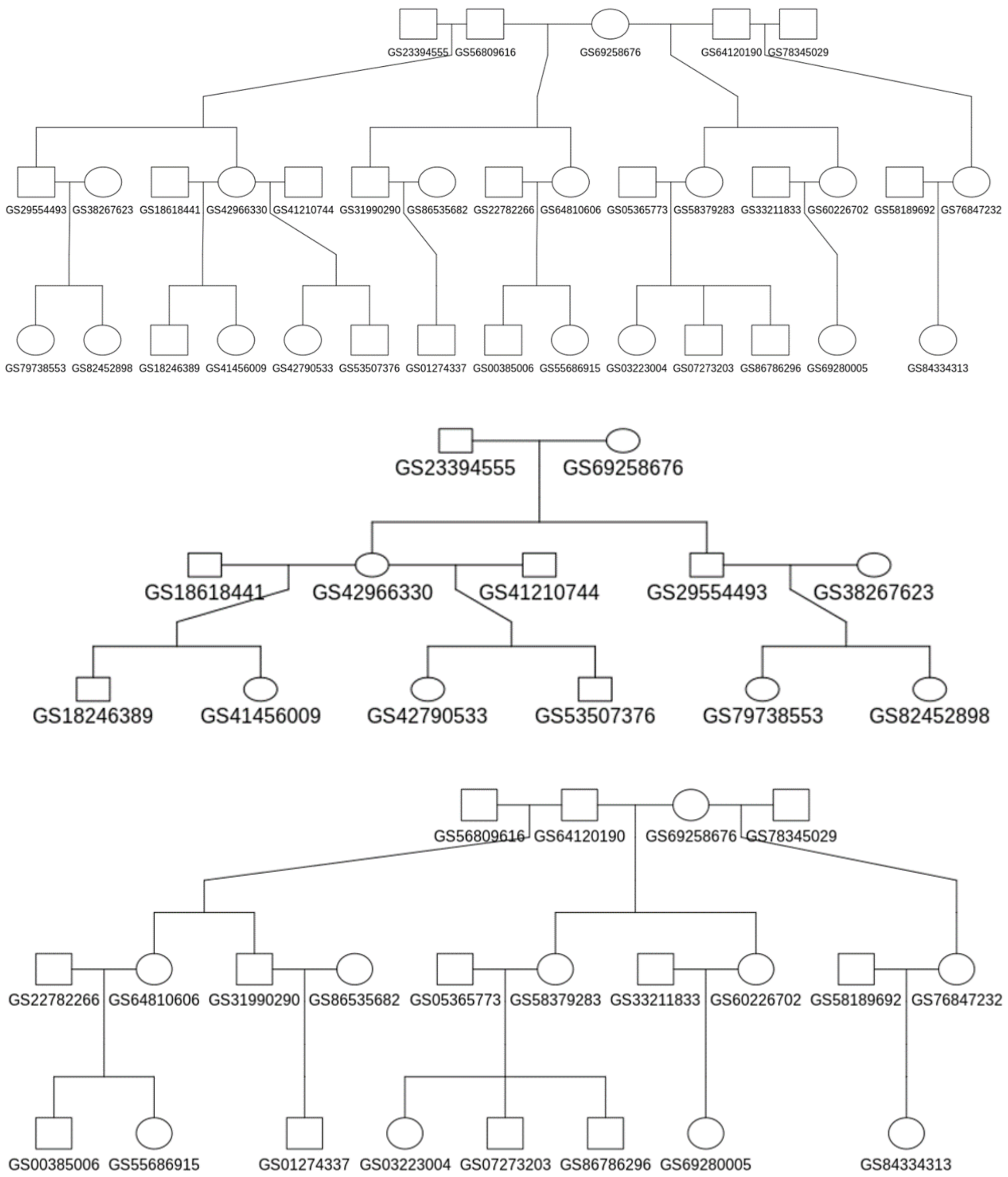


**Supplementary Figure S1:** Pedigree illustrating family structure before (top) and after splitting into two subfamilies based on kinship information. After splitting. the subfamilies are balanced with approximately half of the children in each subgroup.

| Chr | Merlin_OE | | Shapeit2_OE | | Shapeit2_WGS | | Expected | |
|---|---|---|---|---|---|---|---|---|
| | Number | # per Meiosis | Number | # per Meiosis | Number | # per Meiosis | Number | # per Meiosis |
| 1 | 2978 | 10.09 | 885 | 3.00 | 894 | 3.03 | 840 | 2.85 |
| 2 | 2675 | 9.07 | 799 | 2.71 | 794 | 2.69 | 810 | 2.75 |
| 3 | 2398 | 8.13 | 682 | 2.31 | 667 | 2.26 | 687 | 2.33 |
| 4 | 1801 | 6.11 | 634 | 2.15 | 650 | 2.20 | 638 | 2.16 |
| 5 | 2024 | 6.86 | 629 | 2.13 | 626 | 2.12 | 628 | 2.13 |
| 6 | 2199 | 7.45 | 572 | 1.94 | 582 | 1.97 | 583 | 1.98 |
| 7 | 1802 | 6.11 | 555 | 1.88 | 566 | 1.92 | 563 | 1.91 |
| 8 | 1815 | 6.15 | 508 | 1.72 | 487 | 1.65 | 506 | 1.72 |
| 9 | 1594 | 5.40 | 501 | 1.70 | 584 | 1.98 | 497 | 1.68 |
| 10 | 1970 | 6.68 | 521 | 1.77 | 516 | 1.75 | 518 | 1.76 |
| 11 | 1757 | 5.96 | 488 | 1.65 | 510 | 1.73 | 489 | 1.66 |
| 12 | 1634 | 5.54 | 515 | 1.75 | 489 | 1.66 | 496 | 1.68 |
| 13 | 1240 | 4.20 | 362 | 1.23 | 385 | 1.31 | 366 | 1.24 |
| 14 | 1096 | 3.72 | 351 | 1.19 | 360 | 1.22 | 368 | 1.25 |
| 15 | 1007 | 3.41 | 365 | 1.24 | 381 | 1.29 | 375 | 1.27 |
| 16 | 1182 | 4.01 | 407 | 1.38 | 432 | 1.46 | 394 | 1.34 |
| 17 | 911 | 3.09 | 390 | 1.32 | 413 | 1.40 | 409 | 1.39 |
| 18 | 1173 | 3.98 | 366 | 1.24 | 367 | 1.24 | 358 | 1.21 |
| 19 | 770 | 2.61 | 308 | 1.04 | 340 | 1.15 | 303 | 1.03 |
| 20 | 902 | 3.06 | 339 | 1.15 | 339 | 1.15 | 303 | 1.03 |
| 21 | 500 | 1.69 | 184 | 0.62 | 217 | 0.74 | 188 | 0.64 |
| 22 | 484 | 1.64 | 204 | 0.69 | 247 | 0.84 | 205 | 0.69 |
| **ALL** | **33 912** | **114.96** | **10 565** | **35.81** | **10 846** | **36.77** | **10 524** | **35.67** |

**Supplementary Table S2: Comparative analysis of recombination events across autosomes.** This table presents a detailed comparison of the number of recombination events and the recombination events per meiosis across different chromosomes. The data includes results from Merlin_OE (OE: OmniExpress genotype). Shapeit2_OE. Shapeit2_WGS. and the expected values. The 'Merlin_OE' dataset represents OmniExpress genotype data processed using the Merlin software package. while 'Shapeit2_OE' represents data processed with the Shapeit2-duoHMM software package applied to OmniExpress data. and 'Shapeit2_WGS' represents data processed with the Shapeit2-duoHMM software package applied to WGS data. The final row summarizes the total recombination events and their average rates per meiosis.

| Variable | N | min | Q1 | median | mean | Q3 | max |
|---|---|---|---|---|---|---|---|
| **Merlin_OE** | 32 912 | 90 | 18 092 | 44 310 | 143 811 | 110 175 | 88 206 628 |
| **SHapeit2_OE** | 10 565 | 174 | 20 756 | 49 458 | 168 356 | 115 998 | 88 206 628 |
| **SHapeit2_WGS** | 10 846 | 30 | 1 871 | 4 530 | 49 035 | 15 094 | 21 688 832 |

**Supplementary Table S3: the distribution of recombination event interval lengths across chromosomes.** The 'Merlin_OE' dataset represents OmniExpress genotype data processed using the Merlin software. while 'Shapeit2_OE' represents data processed with the Shapeit2-duoHMM software applied to OmniExpress data. and 'Shapeit2_WGS' represents data processed with the Shapeit2-duoHMM software applied to WGS data.

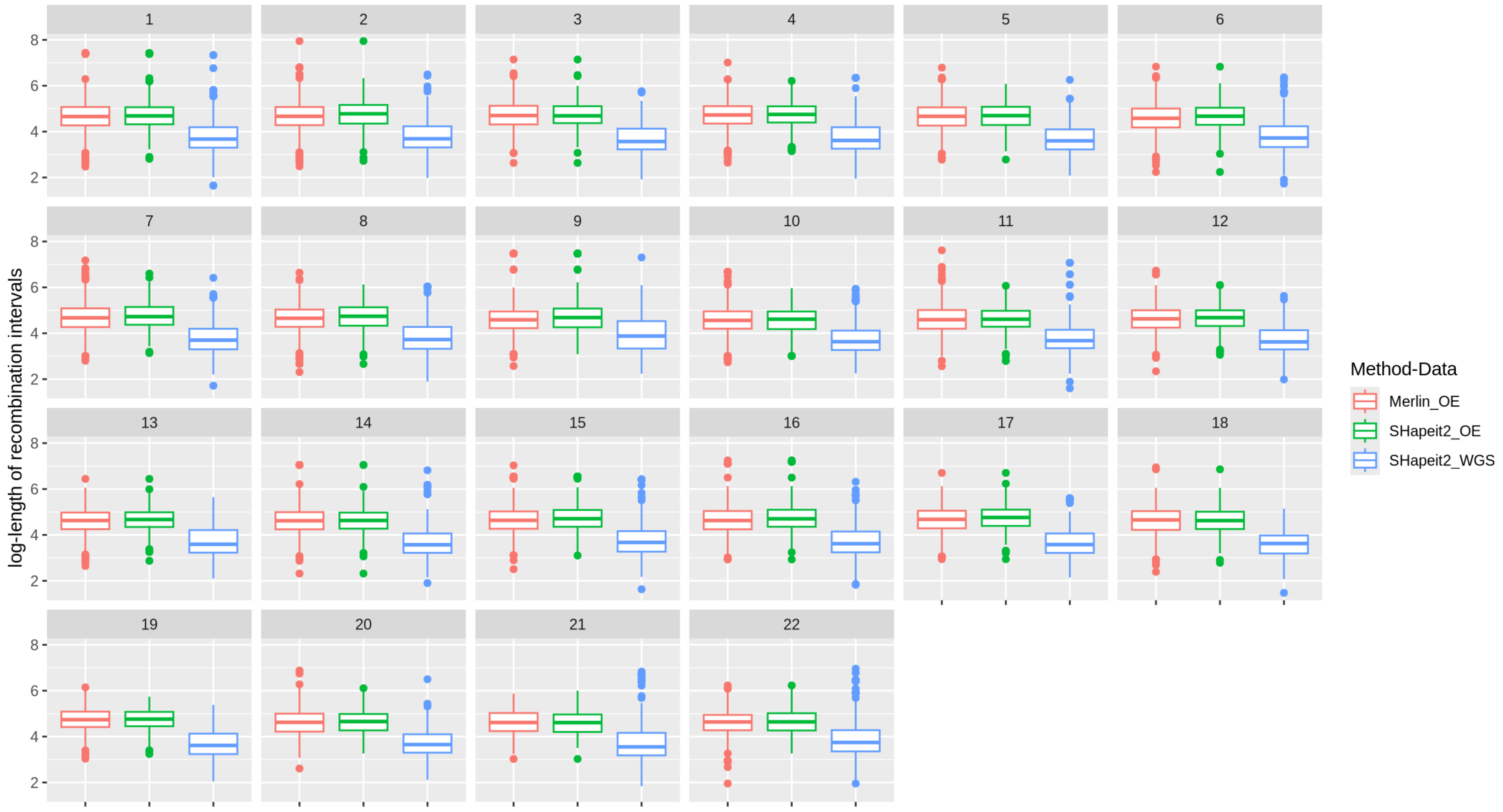


**Supplementary Figure S2: the distribution of recombination event interval lengths across chromosomes.** The 'Merlin_OE' dataset represents OmniExpress genotype data processed using the Merlin software package. while 'Shapeit2_OE' represents data processed with the Shapeit2-duoHMM software package applied to OmniExpress data. and 'Shapeit2_WGS' represents data processed with the Shapeit2-duoHMM software package applied to WGS data.

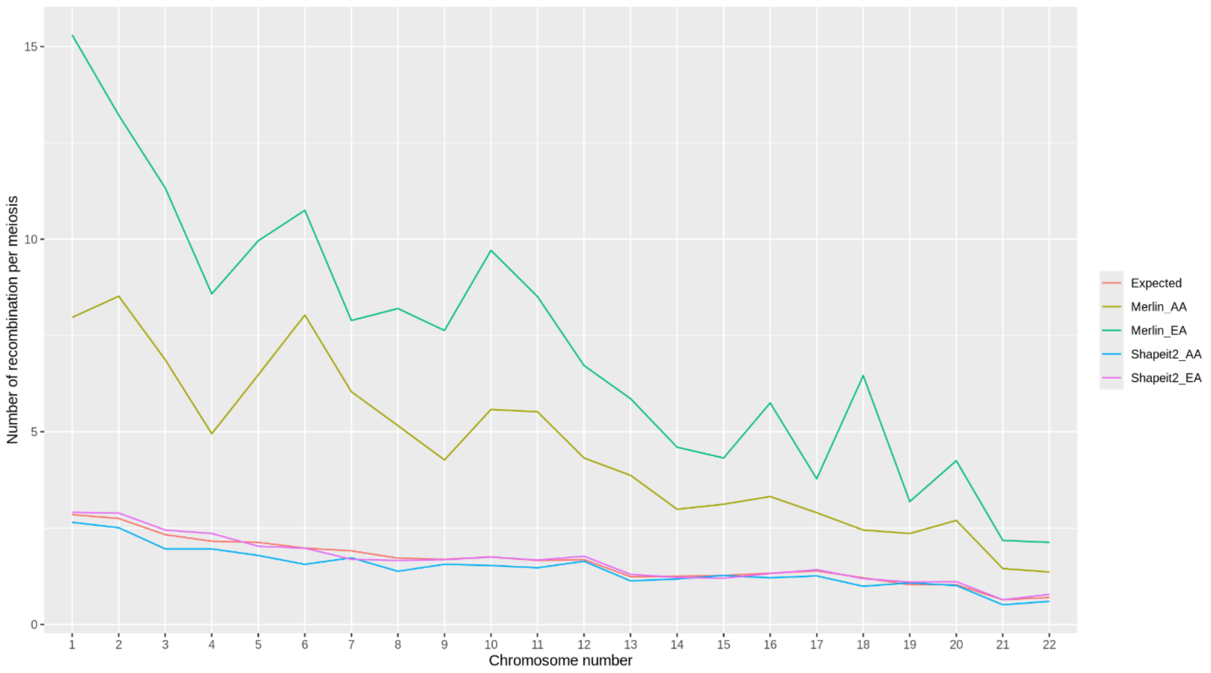


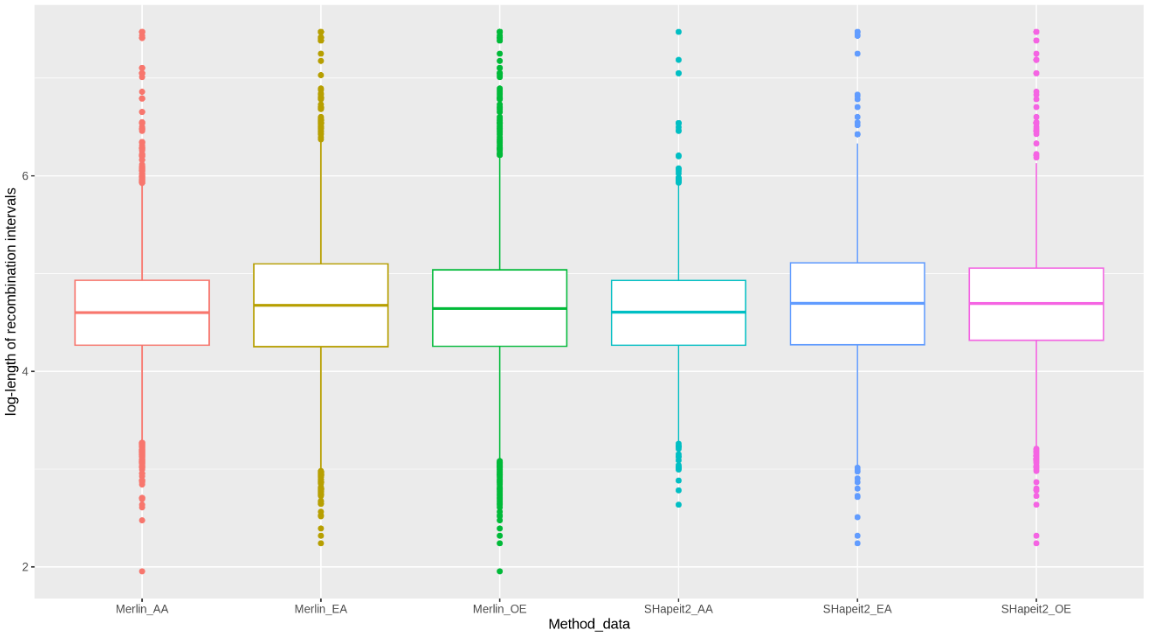


**Supplementary Figure S3: Comparative analysis of recombination events across ethnic groups**. (top) Distribution of recombination events per meiosis. 'Shapeit2_AA' and 'Shapeit2_EA' denote African-American (AA) and European-American (EA) group data processed with the Shapeit2-duoHMM software package on OmniExpress SNP array data. Similarly, 'Merlin_AA' and 'Merlin_EA' represent the corresponding group data processed using the Merlin software package. (bottom) Distribution of recombination event interval lengths across chromosomes, presented on a logarithmic scale.

```
FAMILY GSF0017 [Most Likely]
GS17196750 (FOUNDER) A A A A A A A A A A A A A A A A A A A A A A A A A A A A
GS17196750 (FOUNDER) B B B B B B B B B B B B B B B B B B B B B B B B B B B B
GS35988176 (FOUNDER) C C C C C C C C C C C C C C C C C C C C C C C C C C C C
GS35988176 (FOUNDER) D D D D D D D D D D D D D D D D D D D D D D D D D D D D
GS87227925 (FOUNDER) E E E E E E E E E E E E E E E E E E E E E E E E E E E E
GS87227925 (FOUNDER) F F F F F F F F F F F F F F F F F F F F F F F F F F F F
GS95419745 (FOUNDER) G G G G G G G G G G G G G G G G G G G G G G G G G G G G
GS95419745 (FOUNDER) H H H H H H H H H H H H H H H H H H H H H H H H H H H H
GS52944490 (MATERNAL) E E E E E E E E E E E E E E E E E E E E E E E E E E E E
GS52944490 (PATERNAL) C C C C C C C C C C C C C C C C C C C C C C C C C C C D
GS42295563 (MATERNAL) G G G G G G G G G G G G G G G G G G G G G G G G G G G G
GS42295563 (PATERNAL) E E E E E E E E E E E C E E C E E E E E E E E E E E C D
GS38081276 (MATERNAL) E E E E F F E E E E E E E E E E E E E E E E F F F F F F
GS38081276 (PATERNAL) C C C C C C C C C C C C C C C C C C C C C C C C C C C C
GS94648797 (MATERNAL) A A A A A A A B B B B B B B B B B B B B B B B B B B B B
GS94648797 (PATERNAL) E E E E F F E E E E C C E E E E E E C C C C C C F F F F
GS76032436 (MATERNAL) B B B B B B B B B A A A A A A A A A A B A A A B B B B B
GS76032436 (PATERNAL) C C C C C C C C C C C C C E C C E E E E E C C C C C C C
GS41643637 (MATERNAL) A A A A A A A A B A A A A A A B B B B B B B B B B A A A
GS41643637 (PATERNAL) C C C C C C C C C C C C E C C C C E E E E E F F F F F F
```

**Supplementary Figure S4:** This figure presents an illustrative example of a flow file output generated by MERLIN's haplotyping module in ***--best*** mode. showing the most probable transmission of founder haplotypes within a pedigree (GSF0017 in this case). Each individual is annotated with maternal and paternal haplotypes. inferred from founder alleles under a no-recombination assumption. The flow file encodes allele inheritance across loci. enabling clear visualization of descent patterns.
